# Nitrogen Management on Improving Resilience of Flood-Tolerant (Sub1) Rice Varieties in Flood-Prone Rainfed Lowlands

**DOI:** 10.1101/2024.01.05.574436

**Authors:** A. K. Singh, A. K. Pandey, Ankit Singh, Deeksha Tiwari, Bijendra Singh

## Abstract

Flash flood causing submergence adversely affects rice production in vast areas of rainfed lowlands of South and Southeast Asia. Introgression at the Sub1 locus by molecularly assisted backcrossing incorporated tolerance into varieties traditional/modern high-yielding varieties and has released some varieties for commercial planting by Indian farmers. In the present study, we investigated the application of fractional N to further improve the survival and productivity potential of two Sub1 introgression varieties, Sambha Mahsuri-Sub1, and BR11-Sub1.Thirty days (d) old seedlings were transplanted, and 28-d after transplanting i.e. 58 days old seedlings were completely submerged for 18-d. Full doses of phosphorus and potassium were applied as a basal, while nitrogen was applied in split doses. Plant survival was recorded at 0, 5 and 20 days after de-submergence (AS) to study the recovery dynamics of cultivar Sub1. Results evidently established that varieties with higher biomass, tiller number, total chlorophyll and soluble sugars concentrations before flooding have higher survival rates after water drainage. They also had faster growth and better recovery, which was reflected in yield characteristics and grain yield. The 20-day and 40- day AS N application replenished flood-disturbed soil N pools, resulting in higher N uptake and N-use efficiency. These results may contribute to better nitrogen fertilization programs for rice crops and improve stress tolerance in newly developed tolerant rice varieties. It is evident from perusal of results that treatment T_2_ i.e.1/4 N applied as a basal and rest of residual N applied in three split doses respectively after de-submergence on days 5, 20 and 40 resulted in higher survival thanT1 and T3 respectively minimising mortality rate. This method of nitrogen fertilization (T2) also significantly affected plant^-1^ stem biomass, plant^-1^ tiller number, total chlorophyll, and soluble sugar concentrations before and after flooding. Based on these findings we proposed that application of lower dose of N (30kg/ha) as basal and rest amount of N in three split doses along with P and K (40kg/ha) in the field might be exploited to improve submergence tolerance and to obtain higher yield under flood prone ecosystem due to higher survival after de submergence corresponding to less post oxidative damage through proper N management during before and post submergence period.

## INTRODUCTION

Rice is the predominant staple crop in India and holds a significant position in the country’s food supply, constituting approximately 23% of the overall cultivated land. In India, there are around 16.1 million hectares of rice fields situated in rainfed lowlands, with 44,000 hectares being particularly susceptible to flooding (Haefele and Hijmans, 2007). Flooding is a significant abiotic stressor in numerous lowland rainfed habitats. The number of locations prone to flooding is progressively growing annually, most likely as a result of escalating air temperatures causing intense rainfall and tropical cyclones throughout a significant portion of Asia and Southeast Asia (Easterlings *et al*., 2007; Zeigler and Barcley, 2008). A brief total flood can happen at any point throughout the season, whereas a standing flood typically takes place in the middle of the season and endures for over a month in the field.

Considerable advancements have been achieved in the advancement of marker-assisted backcrossing (MABC) systems for the prominent quantitative trait (QTL) locus SUBMERGENCE1 (SUB1), which is linked to over 70% of the flood tolerance of the rice crop. The MABC technique was used to transfer a significant quantitative trait locus (QTL) called SUB1 onto chromosome 9 of the ‘Swarna’ cultivar. ‘Swarna’ is a popular tall cultivar commonly cultivated in flood-prone areas of South Asia. Consequently, the QTLs were introduced into six specific big Asian rice cultivars that had desirable agronomic and qualitative characteristics favored by farmers. These cultivars are Swarna, Sambha Mahsuri, IR64, BR11, Thadokkam 1 (TDK1), and CR1009 (Mackill *et al*., 2006). Recently, researchers have successfully introduced the SUB1 gene into two different cultivars, PSB Rc18 and Ciherang (Mackill *et al*., 2012; Ismail *et al*., 2013).The field testing of these SUB1 lines shown a notable increase in productivity compared to their recurrent parents when subjected to flooding for a period of 12-18 days during the vegetative phase (Nugraha *et al*., 2013).

Flooding is an abiotic stress that leads to significant decreases in rice yield in Southeast Asia and poses a danger to global food security (FAO, 2016; Yin *et al*., 2017). Flooding can be classified into two categories: water logging and submergence. Water logging refers to the presence of an excessive amount of water in the root zone, causing the soil surface to be covered with water and potentially submerging all or part of the plant (Nishiuchi *et al*., 2012; Sasidhran *et al*., 2017). Short-term total inundation, also known as a flash flood, is the most prevalent and detrimental form of submergence that can happen while rice is in its vegetative development stage. Typically lasting less than 2 weeks, this type of submergence poses significant harm to the crop. The productivity of rainfed lowland rice is consistently hindered by this form of inundation (Iftekharuddaula *et al*., 2016; Pradhan *et al*., 2019; Gautam *et al*., 2019), and can lead to significant harm and death of plants if it persists for more than a week. Asia experiences submergence in almost 22 million hectares of rainfed lowland rice (Gautam *et al*., 2014a). The severity of damage to rice crops due to complete submergence during their vegetative stage is influenced by environmental factors such as elevated temperatures, increased water turbidity, and reduced sun radiation. These variables exacerbate the stress on the crops (Das *et al*., 2009; Ye *et al*., 2018a, b). An energy deficit produced by lack of oxygen can result in decreased absorption of nutrients by plants, leading to the halt of root growth and significant damage and death of the root tips (Bui *et al*., 2019; Liu *et al*., 2015; Singh *et al*., 2014). Furthermore, the plant hormone ethylene builds up in submergence conditions due to its slower diffusive escape compared to the synthesis generated by floods. Multiple studies have documented that ethylene stimulates the elongation of internodes, degradation of chlorophyll, and senescence of leaves. This leads to a decrease in the process of photosynthetic carbon fixation both during and after submergence. These findings have been reported by Ella *et al*. (2003), Jackson (2008), and Yin *et al*. (2017). The overconsumption of energy for shoot elongation and the decrease in carbon fixing during submergence hasten the depletion of carbohydrates. This results in higher mortality rates of submerged plants (Das *et al*., 2005; Bui *et al*., 2019; Ye *et al*., 2018b). Furthermore, studies have shown that the ability of plants to survive submergence is directly related to the retention of chlorophyll and non-structural carbohydrates (Gautam *et al*., 2014a). It is advantageous for plants to limit elongation during flash floods, as stem or internode elongation can compete with vital energy-consuming processes, thereby reducing the likelihood of survival (Bui *et al*., 2019; Ram *et al*., 2002). Additionally, Das *et al*. (2005) found a negative correlation between stem elongation and seedling survival.

Nitrogen is widely recognized as the primary ingredient for enhancing rice yields across various agro-ecosystems (Fageria and Santos, 2014). Nitrogen is crucial for the growth and development of crops. Implementing effective nitrogen management practices both before and after flooding can enhance the survival and productivity of rice. Nitrogen has the potential to enhance the concentration of nitrogen in leaves, the rate of photosynthesis, the delay in leaf aging, and the increase in the quantity of dry matter for grain filling. Consequently, it can enhance the productivity of rice (Hasegawa *et al*., 1994). In addition, N plays a role in enhancing the size of the panicle by increasing the number of filled grains per panicle and increasing grain weight. It also helps to decrease spikelet sterility (Fageria, 2009). Therefore, the optimal utilization of nitrogen (N) can enhance crop productivity by enhancing the number of panicles, increasing the weight of individual grains, and decreasing the percentage of sterility (Fageria, 2007). Furthermore, N has been documented to significantly impact the filling of caryopsis (Juan *et al*., 2006) and the accumulation of dry matter in rice (Mnzava, 2002). The application of nitrogen fertilizer also enhances root development, water utilization, and the uptake of other essential nutrients (Fageria and Santos, 2014). The application of nitrogen fertilizer in rice greatly increased the uptake of nitrogen by grain and straw, as well as the nitrogen use efficiency (Hassan *et al*., 2009). Therefore, it is imperative to use a proper nitrogen management approach that will enhance grain fertility, decrease spikelets sterility, and enhance rice grain yield in locations prone to tidal flooding.

Typically, farmers in flood-prone areas primarily utilize nitrogenous fertilizer and a limited quantity of phosphorus and potassium sources (FAO/WFP, 2016; Soni and Soe, 2016). Within an ecosystem prone to floods, farmers often administer a full dose of nitrogen fertilizer shortly after transplanting, but refrain from applying any nitrogen fertilizer after the flooding occurs. Nevertheless, applying high rates of nitrogen before flooding results in the growth of “succulent” leaves, which have a detrimental effect on plant survival. Conversely, applying nitrogen after flooding promotes regeneration and survival, ultimately enhancing the plant’s ability to tolerate submersion. Hence, the utilization of N-split treatments might effectively mitigate harm and enhance recuperation following partial or complete immersion. Submersion causes leaves to frequently detach and necessitates the growth of new leaves. Sub1 rice cultivars have a high demand for nitrogen (N) in order to expedite recovery during flooding. However, a significant portion of the applied N may be lost as a result of the flooding. It can be inferred that the current suggestions are inadequate for addressing the requirements of flooded crops in general, and specifically the new Sub1 germplasm. Gribaldi *et al*. (2017) reported that applying 50% of the nitrogen fertilizer at planting and the remaining 50% at 42 days following planting resulted in improved crop performance and increased yield. It has been observed that applying more potassium and nitrogen after submergence can enhance the survival and yield of the plants (Dwivedi *et al*., 2018). This study aimed to examine the impact of flooding parameters, including light intensity, temperature, and O_2_ concentration, on the survival of SUB1 rice genotypes. Additionally, we investigated the effects of the timing of N-split application (before and after de-submergence) on plant survival, recovery, and yield after 18 days of complete immersion. The research seeks to enhance efficient nitrogen management in actual field settings at the designated location and will contribute to the augmentation of rice yields and the exploitation of the production capacity of flood-affected regions through the utilization of flood-tolerant cultivars.

## MATERIAL AND METHODS

### Site description

Experiment was carried out in the *Kharif* (monsoon) seasons of 2021-22 and 2022-23 at the Crop Physiology Farm, Acharya Narendra Dev University of Agriculture and Technology, Kumarganj, Ayodhya, India, geo located at 26^0^47’ North and 82^0^12’ East, and an altitude of 113 meters above sea level in the Gangetic alluvium of eastern Uttar Pradesh. The site lies in the humid subtropics, receiving 80% of the total precipitation during the monsoon season (July to end of September) with few showers in winter (Figure 1a and b). The pooled nursery soil test results at the experimental site were as follows: Sand 35%, Silt 49%, Clay 16%, field-capacity 40%,bulk density 1.34 gcm^-2^, pH 7.6, EC 0.23dSm^-1^, organic carbon 0.32%, available N58 ppm, available P_2_O_5_7.5ppm and available K_2_O 235ppm.

**Figure 1:**
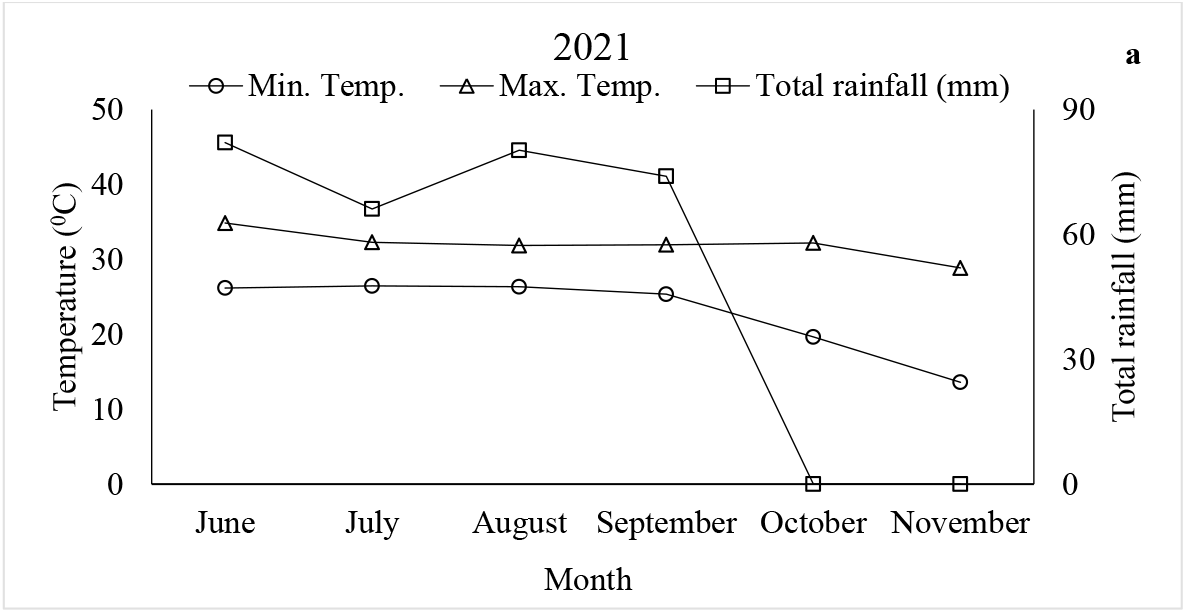
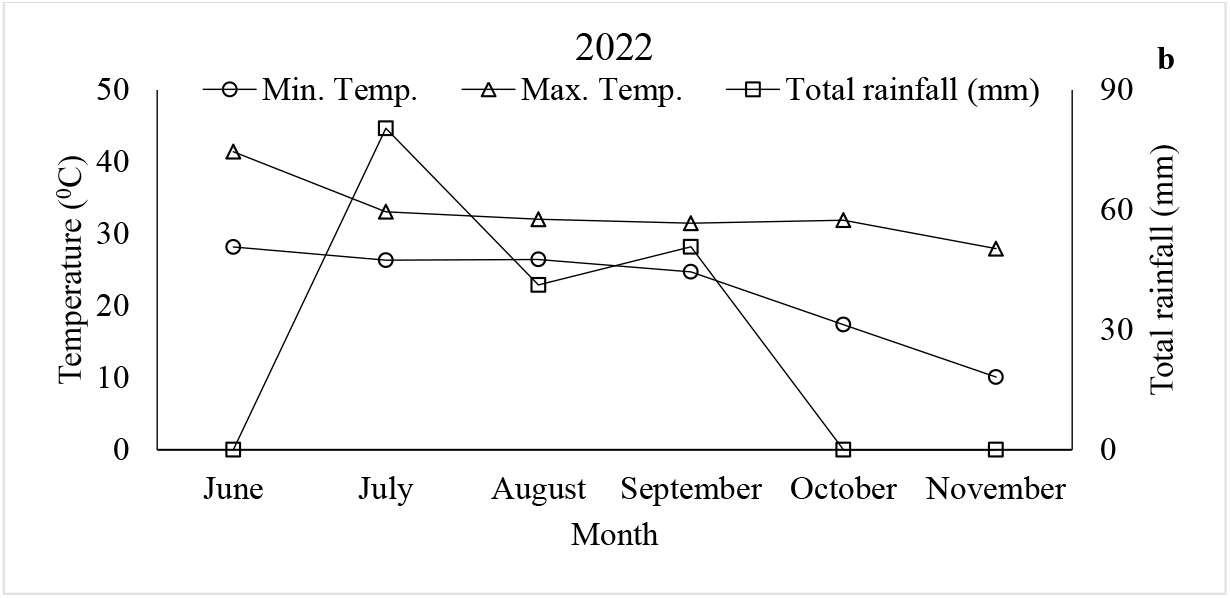
Rainfall and temperature (Maximum and Minimum) during the crop growing periods in 2021 (a) and 2022 (b) at Acharya Narendra Deva University of Agriculture and Technology, Kumarganj, India.

### Experimental layout and crop management

The field experiments were conducted in the submergence pond with a randomized complete block design with 3 replications. The treatment factors were two different SUB1 rice varieties (V) (V_1_: Sambha Mahsuri-Sub1 and V_2_: BR11-Sub1) and three N split doses (T) as depicted in Table 1.

**Table 1:** Schedule of split doses of N applied in the experiment.

| Treatment | Basal (BT) | 5<sup>th</sup> day AS | 20<sup>th</sup> day AS | 40<sup>th</sup> day AS |
| --- | --- | --- | --- | --- |
| T<sub>1</sub> | ½ N | - | ¼ N | ¼ N |
| T<sub>2</sub> | ¼ N | ¼ N | ¼ N | ¼ N |
| T<sub>3</sub> | - | ½ N | ¼ N | ¼ N |

BT: Before transplanting, AS: After de-submergence

**Table 2:**
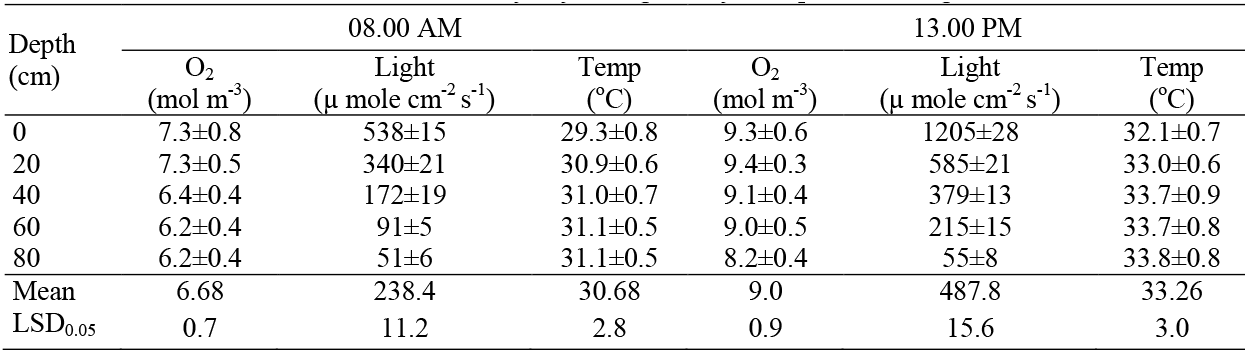
Environmental characterization of flood water in submergence tank during experimental period; Values are means of measurements made every day during 18 days complete submergence ± SEm values.

Well sprouted, healthy and sterilized seeds of both varieties were sown in the nursery according to farmers’ practice, using 100 g m^-2^, the wet method and a 1 m^2^ plot. The equivalent of 2.5 t farm yard manure (FYM) ha^-1^was applied on the nursery plots one week before seeding. In total, 40-40-40 kg N-P_2_O_5_-K_2_Oha^-1^were applied through urea, super phosphate and muriate of potash, respectively. N was applied in two equal split doses: ½ the amount before seeding and the remaining 15-d after seeding (according to farmer practices of the flood prone area), whereas P and K were broadcasted 10-d after seeding.

30-dold seedlings were then transplanted into the submergence pond(size: 20 x 17 x 1.5 m^3^).Seedlings were transplanted at 15 x 15 cm spacing using 3seedlings per hill in plots sized 2.5 x 2 m^2^ during both years. Applied were 120-40-40 kg N-P_2_O_5_-K_2_O ha^-1^using urea, single super phosphate and muriate of potash. The full rates of P and K were applied basal whereas N was applied in split doses as shown in Table 1. At 28-d after transplanting, i.e. 58 days old seedlings were completely submerged for 18-d with turbid water of the nearby water body. 75-80 cm water depth was maintained throughout the period of submergence. Survival (% of all plants in a plot) was determined just after de-submergence (0-d-),5-d-and 20-d after de-submergence. Plant samples were collected before submergence and just-after de-submergence for various measurements. During the experimental period, algal cover was minimized by removing algae from the water surface.

### Flood water quality parameters

Photosynthetic active radiation (PAR, µmol m^-2^ s^-1^), water temperature (°C) and dissolved oxygen (mol m^-3^) was observed at 0, 20, 40, 60 and 80 cm of water depth. All the flood water quality parameters were recorded twice in a day, at 8.00 am and 1.00 pm. PAR was measured using an underwater quantum sensor (Model LI-185 B, Licor Inc., Lincoln, Nebraska) whereas dissolved O_2_ and temperature were measured using a Syland O_2_ electrode as described by Setter *et al*., 1987.

### Assessment of underwater shoot elongation, plant survival, phenology, yield and yield attributes

Plant height, shoot biomass and tiller number hill^-1^ were recorded just before submergence (BS), just after de-submergence (AS) and 20-d after de-submergence (AR, at recovery). Elongation of the shoot was determined by subtracting plant height before submergence from that after de-submergence and expressing it as percentage of plant height before submergence. Underwater shoot elongation day^-1^ was calculated by dividing total increase in plant height by total number of days of submergence. Plant survival was determined by counting the number of hills with green tillers at 0-d, the number of plants that were able to produce at least one new leaf after 5-d, and the number of hills able to regenerate at least one tiller after 20-d of de-submergence. Results were expressed as percentage of hills before submergence for all three measurements.

Crop observations made included phenology indicators and, at maturity, data on yield and yield components *i*.*e*., plant height (cm), days to 50% flowering, days to maturity, ear bearing tillers (EBT), panicle length and weight (cm), 100 seed weight (g), grains panicle^-1^, sterility (%), grain and straw yield (t ha^-1^), and harvest index. Plant height and yield attributes were determined by randomly sampling 10 hills from each plot. Days to 50% flowering occurred when 50% of the hills in each treatment had at least one tiller that reached anthesis. Days to maturity were determined when 95% of the spikelets within each treatment had turned yellow. Panicles were hand-threshed and the fertile and sterile grains were separated by submerging threshed grains in 10% saline water. The samples were oven dried at 70°C to get constant weight. Grain and straw yield were determined on a 10 m^2^ area marked in the middle of each plot. Grains were harvested, dried, and weighed. HI was calculated using the formula of Donald & Hamblin (1976).

### Chlorophyll and total soluble sugar concentrations

Total chlorophyll content was measured with calorimetrically following the procedure of Arnon (1949). 200 mg fresh leaf tissue were homogenized with 5 ml of 80% acetone (v/v) and centrifuged at 4000 rpm for 20 minutes. Absorbance was measured at 645 and 663 nm using a spectrophotometer (Model S177, ELICO, double beam spectrophotometer). Total chlorophyll content was calculated according to:

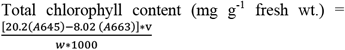

Where v = final volume (ml) and w= final weight of sample (g). Total soluble sugar concentration of the shoot was estimated following the procedure of Yemm & Willis (1954). Shoot samples were dried and grounded to a fine powder and extracted using 80% ethanol (v/v). Extract was then used for soluble sugar analysis after addition of 0.2% anthrone reagent (w/v, in conc. Sulphuric acid), followed by measurement of absorbance at 630 nm using a spectrophotometer (Model S177, ELICO, double beam spectrophotometer). Standard curve was prepared with graded concentrations of glucose.

### Nutrient uptake and N-use efficiency

Soil samples were taken from the plots before start of the experiment and after harvesting of the experiment. Samples were dried and sieved through a 2mm sieve. Subsamples were used to determine the soil pH and EC with a 1:5 soil:water suspension. Nutrients N, P and K were estimated using methods of Lindner (1944) and Jackson (1973), respectively. Nutrient uptake and nitrogen use efficiency were calculated by using the formula:

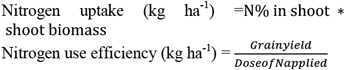

### Statistical analysis

The statistical analysis was carried out for each parameter studied based on the randomized block design following the method of Gomez & Gomez (1984). Associations between parameters were studied using correlation analysis. Means were compared by least significant difference (LSD) at 5% level of significance.

## RESULTS

### Flood water characterization

The flood water temperature during the experimental period at various depths of the pond ranged from 29.3 to 31.1 °C at 8.00 AM and 32.1 to 33.8 °C at 1.00 PM. Slightly higher temperatures were observed at lower depths during both times of measurement; however, differences were statistically non-significant. Photosynthetic active radiation (PAR) was lower during morning hours (50.7 to 538.4 µmol m^-2^ s^-1^) than in the afternoon (54.9 to 1205.4µmol m^-2^ s^-1^) at various water depths of the pond. PAR was reduced by 36.7 to 51.5% when measured at a depth of 20 cm and by 90.5 to 95.5% at the depth of 80 cm, during both measurement times. The dissolved O_2_ content (mol m^-3^) decreased gradually from the surface to lower depths. When measured in the morning, significant differences in O_2_ content were observed only below 20 cm water depth, whereas during the afternoon, differences in O_2_ content were statistically non-significant except at the lowest depth of 80 cm.

### Shoot elongation; shoot biomass, tillers mortality, survival and post-submergence recovery

The nitrogen management treatment T3 (no basal dose of N at transplanting) produced the shortest plants whereas T1 (1/2 basal N dose at transplanting) produced the tallest. However, differences among plant heights were statistically non-significant for both the varieties (Table 3). Submergence caused a significant increase in plant height over their corresponding BS values for all the split N treatments in both varieties with percent elongation ranged from 73.7 to 23.6 and 66.5 to 28.0% for Sambha Mahsuri-Sub1 and BR 11-Sub1, respectively. Across N treatments, BR11-Sub1 had a higher under-water shoot elongation day^-1^ after submergence than Sambha Mahsuri-Sub1 and maintained the shoot elongation rate better with decreasing basal N rate from T1 to T3. At recovery (AR), both the varieties elongated only with about 42% of their values during submergence (AS), and plant height ranged from 101.3 to 109.2 and 101.6 to 110.1 cm for Sambha Mahsuri-Sub1 and BR11-Sub1, respectively (Table 3). Shoot dry biomass BS was not significantly different in all treatments and both varieties (3.15 to 2.87 g plant^-1^). AS, 30.4 to 56.2 and 28.3 to 58.5% reduction in shoot dry biomass was recorded for Sambha Mahsuri-Sub1 and BR11-Sub1, respectively. Maximum reduction was observed in T3 for both varieties. However AR, the highest gain in shoot dry biomass plant^-1^ was noticed in T3 for both varieties (307.9 and 100.0% for Sambha Mahsuri-Sub1 and BR11-Sub1, respectively) and the lowest in T1 (68.0 and 67.2 for Sambha Mahsuri-Sub1 and BR11-Sub1 respectively) (Table 3). Tiller no. plant^-1^ ranged from 13.3 to 10.3 and 13.0 to 9.0 in Sambha Mahsuri-Sub1 and BR11-Sub1, respectively. T1 produced the highest tiller no. plant^-1^ and the lowest was observed in T3. After de-submergence (AS), reductions in tiller no. plant^-1^were recorded in all treatments and both varieties. Higher tiller mortality (41.7 and 41.1% in Sambha Mahsuri-Sub1 and BR11-Sub1, respectively) was noticed in T3 treatment with no basal N dose, whereas least mortality (20 and 18.7% in Sambha Mahsuri-Sub1 and BR11-Sub1, respectively) was recorded in T2 (½ N dose as basal). However, at AR T3 showed a higher recovery in tiller no. plant^-1^ (26.7 in Sambha Mahsuri-Sub1 and 37.7% in BR11-Sub1) whereas other treatments did not significantly influence the tiller regeneration measured 20 d after de-submergence (Table 3).

**Table 3:**
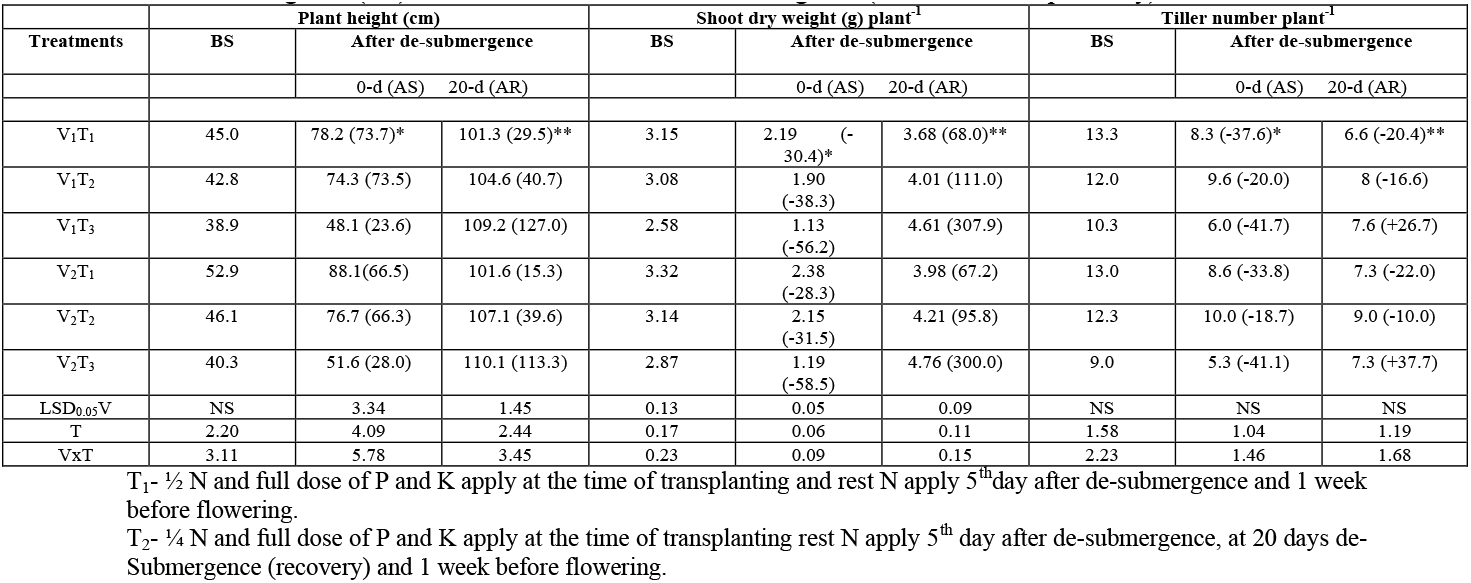
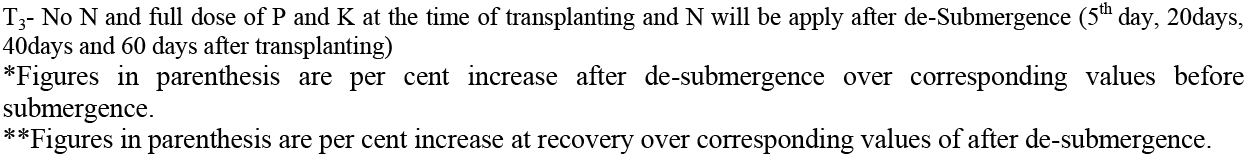
Plant height (cm), Shoot dry weight (g) plant^-1^, Tiller number plant^-1^ of Sambha Mahsuri-Sub1 (V_1_) and BR11-Sub1 (V_2_) as influenced by split N application in the main field (T1 to T3). 58-d old seedlings of both varieties were exposed to 18-d of complete submergence under natural conditions; measurements were recorded before submergence (BS) and at 0- and 20-d after de-submergence (AS and AR, respectively)

| Treatments | Plant height (cm) |  |  | Shoot dry weight (g) plant<sup>-1</sup> |  |  | Tiller number plant<sup>-1</sup> |  |  |
| --- | --- | --- | --- | --- | --- | --- | --- | --- | --- |
|  | BS | After de-submergence |  | BS | After de-submergence |  | BS | After de-submergence |  |
|  |  | 0-d (AS) | 20-d (AR) |  | 0-d (AS) | 20-d (AR) |  | 0-d (AS) | 20-d (AR) |
| V<sub>1</sub>T<sub>1</sub> | 45.0 | 78.2 (73.7)* | 101.3 (29.5)** | 3.15 | 2.19 (-30.4)* | 3.68 (68.0)** | 13.3 | 8.3 (-37.6)* | 6.6 (-20.4)** |
| V<sub>1</sub>T<sub>2</sub> | 42.8 | 74.3 (73.5) | 104.6 (40.7) | 3.08 | 1.90 (-38.3) | 4.01 (111.0) | 12.0 | 9.6 (-20.0) | 8 (-16.6) |
| V<sub>1</sub>T<sub>3</sub> | 38.9 | 48.1 (23.6) | 109.2 (127.0) | 2.58 | 1.13 (-56.2) | 4.61 (307.9) | 10.3 | 6.0 (-41.7) | 7.6 (+26.7) |
| V<sub>2</sub>T<sub>1</sub> | 52.9 | 88.1 (66.5) | 101.6 (15.3) | 3.32 | 2.38 (-28.3) | 3.98 (67.2) | 13.0 | 8.6 (-33.8) | 7.3 (-22.0) |
| V<sub>2</sub>T<sub>2</sub> | 46.1 | 76.7 (66.3) | 107.1 (39.6) | 3.14 | 2.15 (-31.5) | 4.21 (95.8) | 12.3 | 10.0 (-18.7) | 9.0 (-10.0) |
| V<sub>2</sub>T<sub>3</sub> | 40.3 | 51.6 (28.0) | 110.1 (113.3) | 2.87 | 1.19 (-58.5) | 4.76 (300.0) | 9.0 | 5.3 (-41.1) | 7.3 (+37.7) |
| LSD<sub>0.05</sub>V | NS | 3.34 | 1.45 | 0.13 | 0.05 | 0.09 | NS | NS | NS |
| T | 2.20 | 4.09 | 2.44 | 0.17 | 0.06 | 0.11 | 1.58 | 1.04 | 1.19 |
| V×T | 3.11 | 5.78 | 3.45 | 0.23 | 0.09 | 0.15 | 2.23 | 1.46 | 1.68 |

T<sub>1</sub>- ½ N and full dose of P and K apply at the time of transplanting and rest N apply 5<sup>th</sup> day after de-submergence and 1 week before flowering.
T<sub>2</sub>- ¼ N and full dose of P and K apply at the time of transplanting rest N apply 5<sup>th</sup> day after de-submergence, at 20 days de-Submergence (recovery) and 1 week before flowering.
T<sub>3</sub>- No N and full dose of P and K at the time of transplanting and N will be apply after de-Submergence (5<sup>th</sup> day, 20days, 40days and 60 days after transplanting) \*Figures in parenthesis are per cent increase after de-submergence over corresponding values before submergence. \*\*Figures in parenthesis are per cent increase at recovery over corresponding values of after de-submergence.

Submergence reduced plant survival at all stages in both varieties (Table 4). Extent of damage due to submergence ranged from 97-to 100%measured just after de-submergence, from 89.7 to 98.2% at 5-d AS and from 75.8 to 96.7% at 20-d AS. No basal N in the main field (T3) resulted in less survival than treatments with basal N in the main field (T1 & T2). Treatment T2 produced higher survival (98.0 and 98.2% in Sambha Mahsuri-Sub1 and BR11-Sub1, respectively) whereas, least survival was recorded in T3 (89.7 and 92.7% in Sambha Mahsuri-Sub1 and BR11-Sub1, respectively). Sambha Mahsuri-Sub1 showed slightly better survival than BR11-Sub1 at 20-d AS. Treatment T2 (¼^th^ N basal and ¼^th^ N at 5-d AS) resulted in significantly higher survival than the other two treatments in both varieties (Table 4).

**Table 4:** Survival (%) of Sambha Mahsuri-Sub1 (V_1_) and BR11-Sub1 (V_2_) as influenced by split N application in the main field (T1 to T3). 58-d old seedlings of both varieties were exposed to 18-d of complete submergence under natural conditions; measurements were recorded 0-, 5- and 20-days after de-submergence (AS).

| Treatment | Survival % |  |  |
| --- | --- | --- | --- |
|  | After de-submergence |  |  |
|  | 0-d AS | 5-d AS | 20-d AR |
| V<sub>1</sub>T<sub>1</sub> | 99.0 | 94.8 | 93.0 |
| V<sub>1</sub>T<sub>2</sub> | 99.9 | 98.0 | 96.8 |
| V<sub>1</sub>T<sub>3</sub> | 97.1 | 89.7 | 86.7 |
| V<sub>2</sub>T<sub>1</sub> | 98.3 | 96.7 | 92.8 |
| V<sub>2</sub>T<sub>2</sub> | 100.0 | 98.2 | 93.2 |
| V<sub>2</sub>T<sub>3</sub> | 97.7 | 92.7 | 75.8 |
| LSD<sub>0.05</sub> V | 0.05 | 1.34 | 0.94 |
| T | 0.08 | 1.64 | 1.16 |
| VxT | 0.06 | 2.32 | 1.64 |

### Chlorophyll and total soluble sugars concentration

BS Total chlorophyll concentration in leaves increased with increasing basal N rate before transplanting (T1>T2>T3) (Table 5). Total chlorophyll concentration in both varieties decreased significantly during submergence, with highest losses without basal N application (T3; 62.7 and 57.1% for Sambha Mahsuri-Sub1 and BR11-Sub1, respectively) and lowest losses in T1 (42.5 and 44.2% for Sambha Mahsuri-Sub1 and BR11-Sub1, respectively). However, at recovery (20 d) maximum total chlorophyll concentration was recorded in T3 which received 1/2 N dose at 5-d of de-submergence, followed by T2 (1/4^th^ N at 5-d of de-submergence) and least in T1 (no N at 5-d of de-submergence) (Table 5).

**Table 5:**
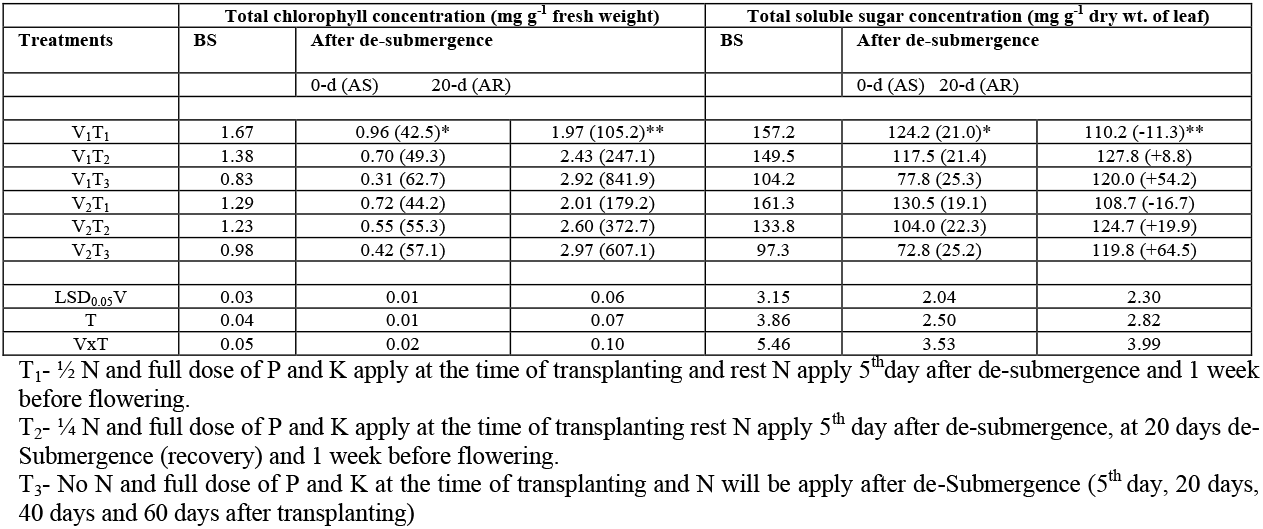
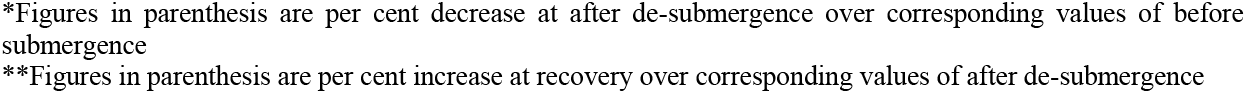
Total chlorophyll concentration (mg g^-1^ fresh weight) and total soluble sugar concentration (mg g^-1^ dry wt. of leaf) of Sambha Mahsuri-Sub1 (V_1_) and BR11-Sub1 (V_2_) as influenced by split N application in main field (T1 to T3). 58-d old seedlings of both varieties were exposed to 18-d of complete submergence under natural conditions; measurements were recorded before submergence (BS) and 0- and 20-d after de-submergence (AS and AR, respectively)

| Treatments | Total chlorophyll concentration (mg g<sup>-1</sup> fresh weight) |  |  | Total soluble sugar concentration (mg g<sup>-1</sup> dry wt. of leaf) |  |  |
| --- | --- | --- | --- | --- | --- | --- |
|  | BS | After de-submergence |  | BS | After de-submergence |  |
|  |  | 0-d (AS) | 20-d (AR) |  | 0-d (AS) | 20-d (AR) |
| V<sub>1</sub>T<sub>1</sub> | 1.67 | 0.96 (42.5)* | 1.97 (105.2)** | 157.2 | 124.2 (21.0)* | 110.2 (-11.3)** |
| V<sub>1</sub>T<sub>2</sub> | 1.38 | 0.70 (49.3) | 2.43 (247.1) | 149.5 | 117.5 (21.4) | 127.8 (+8.8) |
| V<sub>1</sub>T<sub>3</sub> | 0.83 | 0.31 (62.7) | 2.92 (841.9) | 104.2 | 77.8 (25.3) | 120.0 (+54.2) |
| V<sub>2</sub>T<sub>1</sub> | 1.29 | 0.72 (44.2) | 2.01 (179.2) | 161.3 | 130.5 (19.1) | 108.7 (-16.7) |
| V<sub>2</sub>T<sub>2</sub> | 1.23 | 0.55 (55.3) | 2.60 (372.7) | 133.8 | 104.0 (22.3) | 124.7 (+19.9) |
| V<sub>2</sub>T<sub>3</sub> | 0.98 | 0.42 (57.1) | 2.97 (607.1) | 97.3 | 72.8 (25.2) | 119.8 (+64.5) |
| LSD<sub>0.05</sub>V | 0.03 | 0.01 | 0.06 | 3.15 | 2.04 | 2.30 |
| T | 0.04 | 0.01 | 0.07 | 3.86 | 2.50 | 2.82 |
| VxT | 0.05 | 0.02 | 0.10 | 5.46 | 3.53 | 3.99 |

\*Figures in parenthesis are per cent decrease at after de-submergence over corresponding values of before submergence
\*\*Figures in parenthesis are per cent increase at recovery over corresponding values of after de-submergence

Parallel to leaf chlorophyll concentration, significantly lower total pre-submergence soluble sugar was observed in T3, followed by T2 and T1 for both varieties (Table 5). After de-submergence, maximum reduction in total soluble sugars concentration was recorded in T3 (25.3 and 25.2%) followed by T2 (21.4 and 22.3%) and T1 (21.0 and 19.1%) in Sambha Mahsuri Sub1 and BR11-Sub1 respectively. However AR, treatments with a N dose at 5-d after de-submergence (T2 and T3) did increase their total soluble sugar compared to BS values, whereas in T1 (no N at 5-d after de-submergence) total soluble sugar concentration in leaves decreased between AS to BS (Table 5).

### Phenology, yield attributes and yield

Days to 50% flowering was significantly affected by submergence in both varieties. Maximum delay to 50% flowering was recorded in T1 followed by T3 and least in T2 (Figure 2a). However, BR11-Sub1 experienced comparatively less delay in 50% flowering than Sambha Mahsuri-Sub1. The trend in days to maturity was similar to that of days to 50% flowering (Figure 2b). The maximum delay to 50% flowering and the maximum delay to physiological maturity was recorded in treatment T3 for both the Sub1 cultivars.

**Figure 2:**
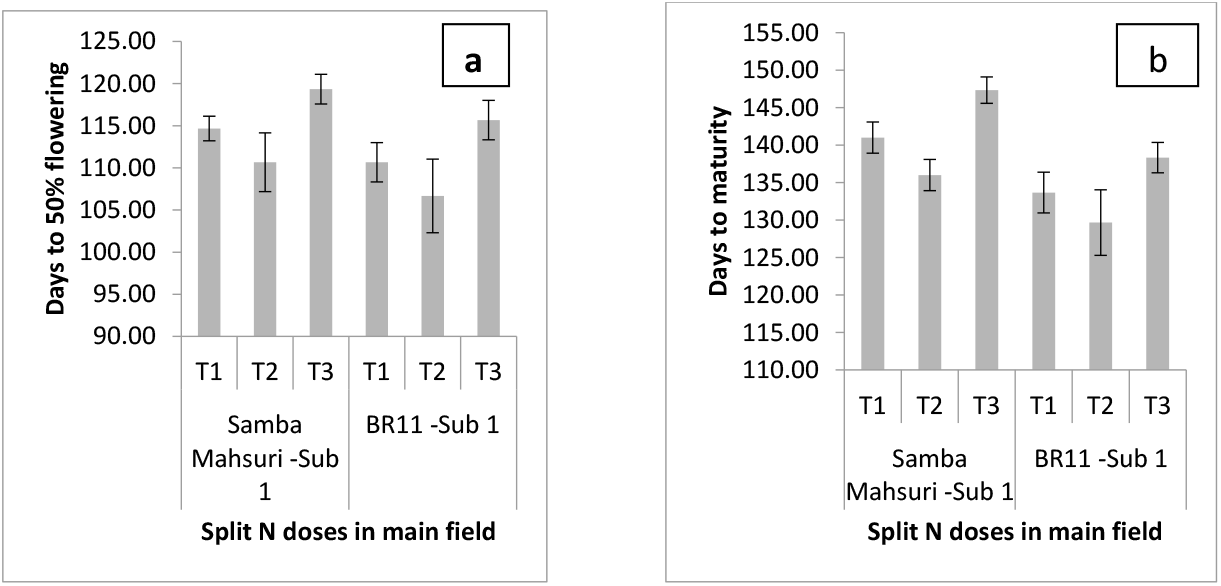
Days to 50% flowering (2a) and days to maturity (2b) of Sambha Mahsuri-Sub1 and BR11-Sub1 subjected to 18-d of complete submergence as influenced by split N doses in the main field; vertical bars in each column represents standard error. T_1_-½ N and full dose of P and K apply at the time of transplanting and rest N apply 5^th^day after de-submergence and 1 week before flowering. T_2_-¼ N and full dose of P and K apply at the time of transplanting rest N apply 5^th^ day after de-submergence, at 20 days de-Submergence (recovery) and 1 week before flowering. T_3_-No N and full dose of P and K at the time of transplanting and N will be apply after de-Submergence (5^th^ day, 20days, 40days and 60 days after transplanting)

At maturity, maximum plant height for both varieties was recorded in T2 followed by T3 and T1; moreover, differences were found statistically significant. Ear bearing tillers plant^-1^ (EBT) ranged from 5.5 to 6.6 and from 5.1 to 6.7 for Sambha Mahsuri-Sub1 and BR11-Sub1, respectively. T2 produced maximum EBT plant^-1^, followed by T1 and was lowest in T3 (Table 6). Comparatively higher panicle length and panicle weight (26.9 to 30.0 cm and 3.92 to 4.73 g) were observed in BR11-Sub1 than in Sambha Mahsuri-Sub1 (22.7 to 24.2 cm and 3.13 to 3.75 g). However, the differences in panicle length of both the Sub 1 cultivars were found significant among all the treatments, and the panicle weight of both Sub1 cultivars was significantly higher in treatment T2 over the other two treatments. The 100 seed weight (g) of BR11-Sub1 was higher for all the treatments than that of Samba Mahsuri-Sub1; also the differences among treatments for both varieties were statistically significant. In contrast to the 100 seed weight, grain no. panicle^-1^ was higher for all the treatments in Sambha Mahsuri-Sub1. In both varieties, T2 recorded higher grains no. panicle^-1^ but differences were statistically non significant. Lower spikelet sterility was observed in BR11-Sub1 than in Sambha Mahsuri-Sub1 (9.1 to 10.9% and 14.6 to 18.0%, respectively). T3 showed higher sterility in both varieties whereas minimum sterility was recorded in T2 (14.6% in SambhaMahsuri-Sub1) and (9.1% in BR11-Sub1) (Table 6).

**Table 6:**
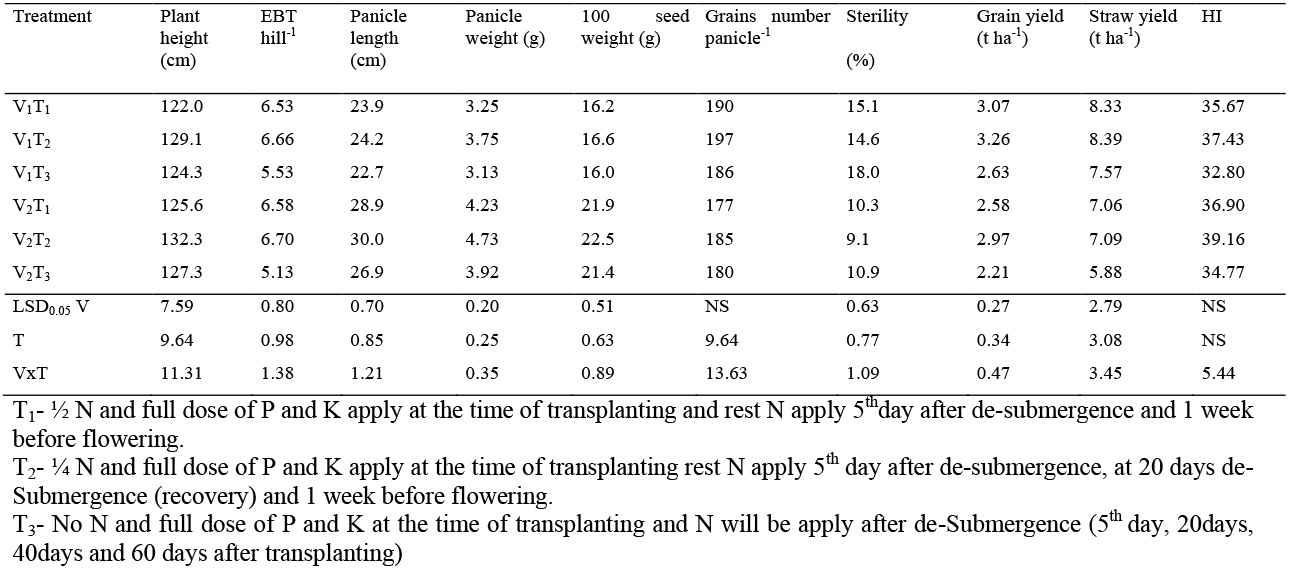
Yield and yield attributes of Sambha Mahsuri-Sub1 (V_1_) and BR11-Sub1 (V_2_) as influenced by split N applications (T1 to T3). 58-d old seedlings of both varieties were exposed to 18-d of complete submergence under natural conditions

Higher grain and straw yield were observed in Sambha Mahsuri-Sub1 (Table 6). Grain yields ranged from 2.63 to 3.26 and 2.21 to 2.97tha^-1^ for Sambha Mahsuri-Sub1 and BR11-Sub1, respectively. T2 produced the highest grain yield. Harvest indices varied from 32.8 to 37.43 and 34.77 to 39.16 for Sambha Mahsuri-Sub1 and BR11-Sub1, respectively moreover statistically analysis of harvest index was found non-significant among individual cultivars and treatments; however the interaction between cultivar and treatment was found significant (Table 6).

### N uptake and N use efficiency

Both Sub1 varieties showed very similar patterns of N uptake (kg ha^-1^). BS, treatments with N application (T1 and T2) recorded higher N uptake than treatments with no N application (T3), and both varieties showed a similar pattern AS (Figure 3). AR, maximum N uptake was measured in T2 (1/4^th^ N applied 5^th^ d after de-submergence) followed by T3 (1/2 N applied 5^th^ d after de-submergence).Therefore, T2 exhibited the highest N use efficiency, followed by T3 and was lowest in T1 for both varieties. Between varieties, BR11-Sub1 showed comparatively lower N use efficiency than Sambha Mahsuri-Sub1 for all N treatments (Figure 3). Among varieties BR 11 Sub 1 showed comparatively lower N uptake as well as N use efficiency than Sambha Mahsuri Sub 1 for all N treatments.

**Figure 3:**
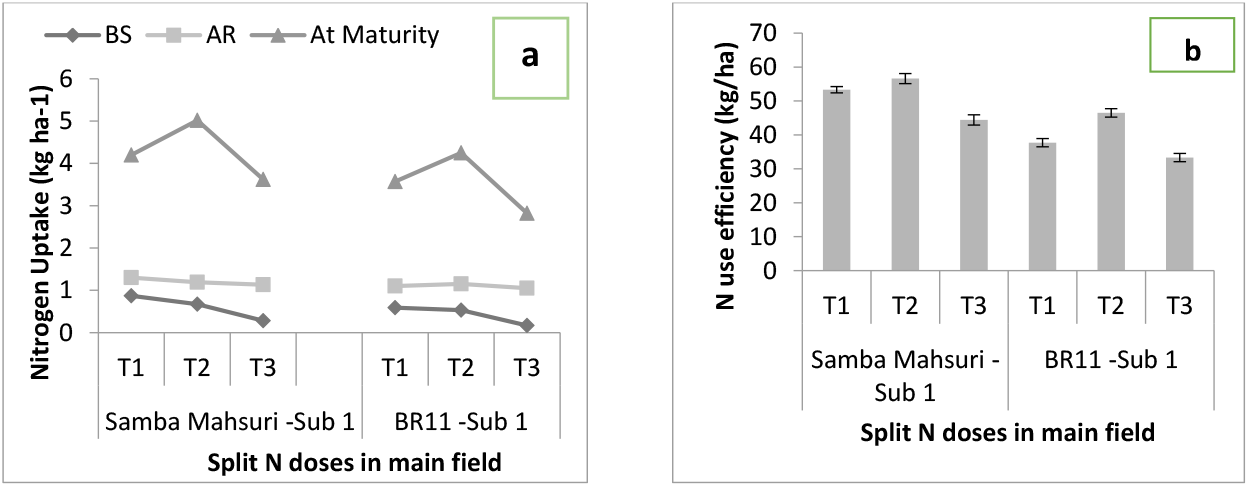
N uptake (kg ha^-1^) 4 (a) and N use efficiency (kg ha^-1^) 4 (b) of Sambha Mahsuri-Sub1 and BR11-Sub1 subjected to 18-d of complete submergence as influenced by split N doses in main field; vertical bars in each line represents standard error. T_1_-½ N and full dose of P and K apply at the time of transplanting and rest N apply 5^th^day after de-submergence and 1 week before flowering. T_2_-¼ N and full dose of P and K apply at the time of transplanting rest N apply 5^th^ day after de-submergence, at 20 days de-Submergence (recovery) and 1 week before flowering. T_3_-No N and full dose of P and K at the time of transplanting and N will be apply after de-Submergence (5^th^ day, 20days, 40days and 60 days after transplanting)

## DISCUSSION

Submergence stress affects over 22 million ha rice area and cause an annual yield loss of over 600 million to 1 billion US Dollar annually in Asian countries. This issue is likely to get aggravated, as the flood-affected areas are anticipated to increase substantially in the era of global warming (Coumore and Ramstrof, 2012).

### Flood water characterization and tolerance to submergence

Submergence is characterized by a series of intricate non-living stresses. The severity of the harm caused by complete submergence mostly relies on the length of time and the parameters of the flood water, namely its temperature, turbidity, and the degree of light penetration (Ram *et al*., 1999, 2002; Das *et al*., 2009). Weather variables, such as air temperature and cloudiness, have a significant impact on most of these elements. During the experimental period of submergence, the concentration of O_2_ was observed to be lower in the morning (8.00 AM) and greater in the afternoon (13.00 PM). During the night, the process of respiration by submerged rice plants and other microorganisms in flood water leads to a decrease in the oxygen levels, resulting in the depletion of oxygen. The process of water oxygenation during daylight hours is primarily attributed to the underwater photosynthesis of rice plants and algae. The diffusion of gases in water is significantly reduced by a factor of 10^-4^ compared to air, as observed by Armstrong in 1979. This limitation hinders the exchange of CO_2_ and O_2_, which are essential for photosynthesis and respiration. Additionally, it leads to an increase in the concentration of the gaseous hormone ethylene within cells. When combined with gibberellic acid, this hormone induces elongation of the stem internode. The decrease in the intensity of light that reaches underwater leaves, particularly in murky water, accelerates the process of chlorosis and leaf senescence (Ella *et al*., 2003). Hence, the primary outcome of submergence is a decrease in photosynthesis and respiration. The temperature of the surface and underwater environments is a crucial determinant of the survival of submerged vegetation. Significantly higher surface and underwater temperatures were detected throughout the afternoon (ranging from 32.1 to 33.8 °C at 13.00 PM) compared to the morning (ranging from 29.3 to 31.1 °C at 8.00 AM). However, there were no significant temperature differences at different depths. Ram *et al*. (2002) and Das *et al*. (2009) found evidence of an inverse relationship between water temperature and survival. Evidently, the survival rate in transparent water declined by 8% for every 1°C rise in temperature beyond 26°C. In rice, the ability of roots to respire is facilitated by the transport of oxygen from the shoot through a specialized tissue called aerenchyma. However, this transport is hindered when the plant is submerged. Under conditions of submergence, the plant hormone ethylene builds up due to its slower rate of diffusive escape compared to the rate of synthesis generated by floods. Multiple studies have documented that ethylene stimulates the growth of internodes, the breakdown of chlorophyll, and the aging of leaves, leading to a decrease in the process of photosynthetic carbon fixation during and after being submerged (Ella *et al*., 2003; Jackson, 2008; Yin *et al*., 2017). The overconsumption of energy for shoot elongation and the decrease in carbon fixing during submergence expedite the depletion of carbohydrates. The mortality of submerged plants is elevated by this phenomenon (Das *et al*., 2005; Bui *et al*., 2019; Ye *et al*., 2018b).

### Response of Sub1 rice varieties to complete submergence

A study conducted in the Eastern regions of India on rainfed lowland rice has discovered two specific physiological characteristics that determine the ability to tolerate submergence. These attributes are: 1. minimum elongation of shoots when submerged underwater, and 2. a high concentration of storage carbohydrates before submergence (Srivastava *et al*., 2007). Recent molecular studies have revealed a single polygenic locus, known as SUB1, on chromosome 9, which plays a crucial role in submergence tolerance in rice. The Sub1 gene possesses alleles that enable the plant to sustain elevated quantities of stored carbohydrates while minimizing shoot elongation in submerged conditions. After resurfacing, the plant resumes the process of leaf growth and the preservation of chlorophyll, as demonstrated in the FR13A submergence-tolerant genotype model (Fukao & Bailey-Serres, 2008).

During our experiment, a period of 18 days of complete submergence resulted in little plant death. The survival rate of the Sub1 variety ranged from 97.1% to 100.0%, as measured immediately after the plants were no longer submerged (Table 4). The survival rates, measured at 5 and 20 days after stress (dAS), which indicate the extent of damage caused by oxidation and subsequent recovery, varied between 89.7% and 98.2% and between 75.8% and 96.8%, respectively, for both Sub1 types (Table 4). Oxidative stress arises when there is an imbalance in the usual plant processes, and the reintroduction of oxygen after a period of “low oxygen” stress caused by complete submergence leads to an increase in the production of reactive oxygen species (ROS). Reactive oxygen species (ROS) are produced through several enzymatic and non-enzymatic processes. The main targets of ROS are membrane lipids, leading to cell death in the end (Szarka *et al*., 2012).

Rice plants have a variety of physiological and metabolic disruptions while submerged, which hinders their growth and development (Ismail 2013; Singh *et al*., 2014b). The study conducted by Gautam *et al*. (2014 a) discovered a positive correlation between plant survival and the retention of chlorophyll and non-structural carbohydrates during submergence. In their study, Singh *et al*.,2014b emphasized the importance of chlorophyll retention in ensuring survival during and after submergence. This is because chlorophyll retention enables underwater photosynthesis and facilitates quicker recovery once the water recedes. Both SUB1 types exhibited a decrease of around 45% in the overall chlorophyll content and a 25% decrease in the total soluble sugar content following de-submergence, as indicated in Table 5. Nevertheless, they managed to enhance the overall chlorophyll content by a factor of 5-6 and recover 20-50% of the total soluble sugars during the restoration phase (Table 5). This clearly demonstrates a rapid ability to regenerate, which ultimately translates into improved yield characteristics and overall crop productivity (Table 6).

After a period of 20 days following de-submergence, the survival rate was shown to be inversely correlated with elongation (-0.47; Table 7). This finding shows that decreased elongation underwater is regarded advantageous for submergence tolerance (Jackson & Ram, 2003). Rice genotypes that exhibit reduced growth and respiration in response to flooding are selected for cultivation in flash-flood prone regions (Bailey-Serres & Voesenek, 2008). The stunted shoot growth observed during flooding is a result of the plant saving its carbohydrate reserves by allocating only the minimum amount necessary for maintenance growth. This strategy allows for an extended energy supply, ensuring enhanced survival. The variations in elongation and survival could be attributed to a heightened sensitivity to ethylene, leading to more pronounced leaf chlorosis during submergence. This, in turn, increases the vulnerability of intolerant genotypes to lodging when the water level recedes (Gautam *et al*., 2014).The reduction in shoot elongation mediated by SUB1 during complete submergence is an adaptation characteristic that aids in minimizing carbohydrate usage, hence enhancing survival (Mackill *et al*., 2012).

**Table 7:**
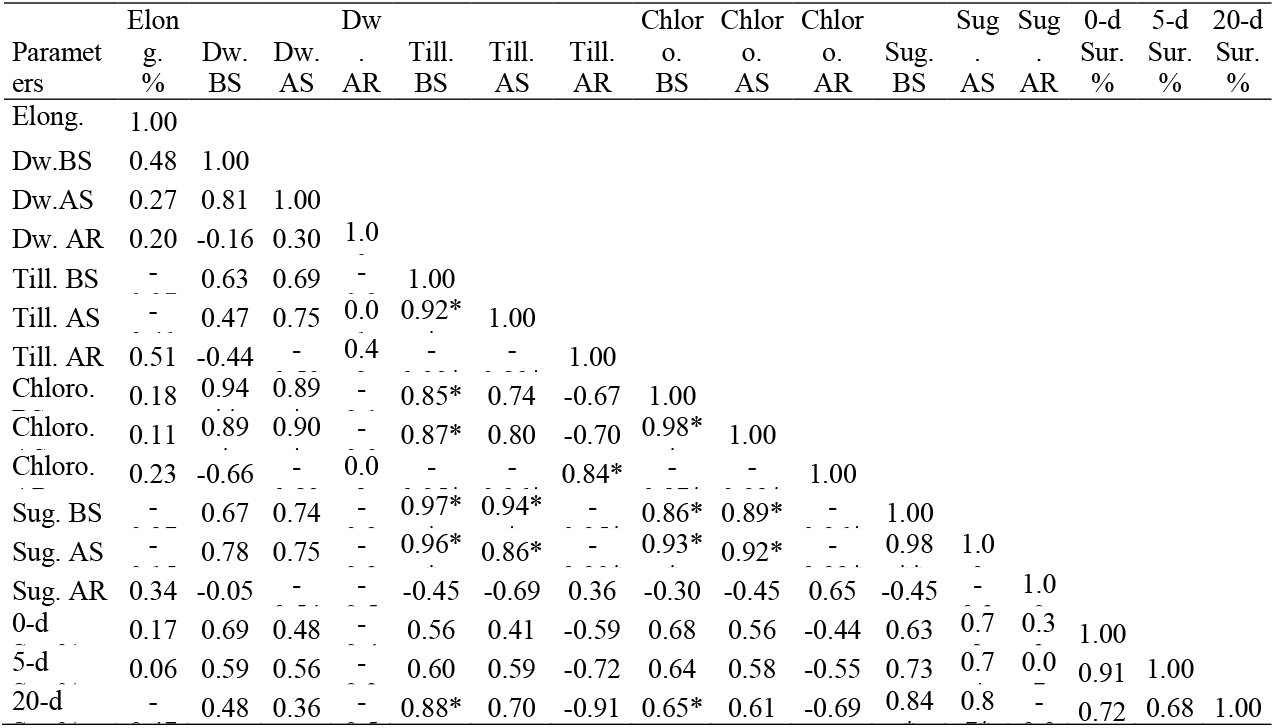
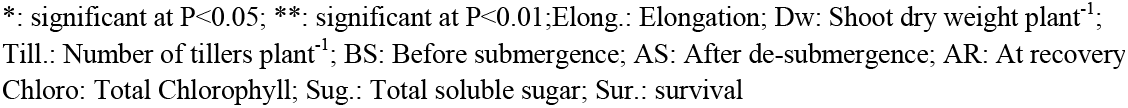
Relationship among survival% and different agronomic and physiological parameters influenced by split application of N in main field; 58-d old seedlings of both varieties were exposed to 18-d of complete submergence under natural conditions

| Parameters | Elon g. % | Dw. BS | Dw. AS | Dw. AR | Till. BS | Till. AS | Till. AR | Chlor. BS | Chlor. AS | Chlor. AR | Sug. BS | Sug. AS | Sug. AR | 0-d % | 5-d % | 20-d % |
| --- | --- | --- | --- | --- | --- | --- | --- | --- | --- | --- | --- | --- | --- | --- | --- | --- |
| Elong. | 1.00 |  |  |  |  |  |  |  |  |  |  |  |  |  |  |  |
| Dw.BS | 0.48 | 1.00 |  |  |  |  |  |  |  |  |  |  |  |  |  |  |
| Dw.AS | 0.27 | 0.81 | 1.00 |  |  |  |  |  |  |  |  |  |  |  |  |  |
| Dw. AR | 0.20 | -0.16 | 0.30 | 1.0 |  |  |  |  |  |  |  |  |  |  |  |  |
| Till. BS | - | 0.63 | 0.69 | - | 1.00 |  |  |  |  |  |  |  |  |  |  |  |
| Till. AS | - | 0.47 | 0.75 | 0.0 | 0.92* | 1.00 |  |  |  |  |  |  |  |  |  |  |
| Till. AR | 0.51 | -0.44 | - | 0.4 | - | - | 1.00 |  |  |  |  |  |  |  |  |  |
| Chloro. | 0.18 | 0.94 | 0.89 | - | 0.85* | 0.74 | -0.67 | 1.00 |  |  |  |  |  |  |  |  |
| Chloro. | 0.11 | 0.89 | 0.90 | - | 0.87* | 0.80 | -0.70 | 0.98* | 1.00 |  |  |  |  |  |  |  |
| Chloro. | 0.23 | -0.66 | - | 0.0 | - | - | 0.84* | - | - | 1.00 |  |  |  |  |  |  |
| Sug. BS | - | 0.67 | 0.74 | - | 0.97* | 0.94* | - | 0.86* | 0.89* | - | 1.00 |  |  |  |  |  |
| Sug. AS | - | 0.78 | 0.75 | - | 0.96* | 0.86* | - | 0.93* | 0.92* | - | 0.98 | 1.0 |  |  |  |  |
| Sug. AR | 0.34 | -0.05 | - | - | -0.45 | -0.69 | 0.36 | -0.30 | -0.45 | 0.65 | -0.45 | - | 1.0 |  |  |  |
| 0-d | 0.17 | 0.69 | 0.48 | - | 0.56 | 0.41 | -0.59 | 0.68 | 0.56 | -0.44 | 0.63 | 0.7 | 0.3 | 1.00 |  |  |
| 5-d | 0.06 | 0.59 | 0.56 | - | 0.60 | 0.59 | -0.72 | 0.64 | 0.58 | -0.55 | 0.73 | 0.7 | 0.0 | 0.91 | 1.00 |  |
| 20-d | - | 0.48 | 0.36 | - | 0.88* | 0.70 | -0.91 | 0.65* | 0.61 | -0.69 | 0.84 | 0.8 | - | 0.72 | 0.68 | 1.00 |

\*: significant at P<0.05; \*\*: significant at P<0.01;Elong.: Elongation; Dw: Shoot dry weight plant<sup>-1</sup>; Till.: Number of tillers plant<sup>-1</sup>; BS: Before submergence; AS: After de-submergence; AR: At recovery; Chloro: Total Chlorophyll; Sug.: Total soluble sugar; Sur.: survival

The data in Table 3 clearly demonstrate a decrease in shoot dry weight per plant before submergence, during submergence, and during the recovery period, indicating the negative impact of submergence on dry weight. Sub1 variety exhibit sufficient dry matter retention, enabling accelerated growth and development following de-submergence. Consequently, they demonstrate superior survival rates compared to genotypes that experience more dry matter depletion during submergence (Chaturvedi *et al*., 1996). Table 7 shows that there is a positive correlation of 0.48 and 0.36 between the 20-day survival data and the pre- and post-submergence shoot dry weight of plant 1, respectively.

The correlation between survival and pre-submergence tiller number per plant was 0.88, while the correlation between survival and post-submergence tiller number per plant was 0.70 (Table 7). However, when submerged, the Sub1 cultivars exhibited a significant decrease in the number of tillers per plant. The decrease in the number of tillers per plant under submergence is caused by the death and decomposition of live components and/or the decreased provision of carbohydrates from photosynthesis occurring underwater. The injection of phosphorus (P) at the base of the plants and the application of nitrogen (N) after flooding resulted in reduced shoot elongation, plant mortality, and ultimately increased plant survival (Gautam *et al*., 2014a). Singh *et al*. (2014a) and Bhowmik *et al*. (2014) found that the application of nitrogen (N) 5-7 days after sowing (DAD) enhanced survival, increased plant population at recovery, and consequently resulted in greater grain output. Singh *et al*. (2014a) found that the increased regrowth of plants following de-submergence was linked to the application of nitrogen after de-submergence, rather than the application before submergence. In a similar vein, Gautam *et al*. (2015) discovered that applying N following de-submergence resulted in an increase in the quantity of green leaves, photosynthetic rate, biomass, and yield.Prior to submergence, the treatments exhibited varying amounts of total chlorophyll content as a result of the varied doses of nitrogen applied. Higher doses of nitrogen treatments resulted in increased total chlorophyll concentrations, as shown in Table 5. A strong positive connection was observed between the content of total chlorophyll in leaves before submergence and survival (0.65*; Table 7). The study found no significant connections between the concentration of total chlorophyll in post-submergence leaves and survival (0.61; Table 7).

This suggests that the concentration of chlorophyll before submergence plays a crucial role in determining survival. In their study, Singh *et al*. (2014b) emphasized the importance of chlorophyll retention for survival during and after submergence. This is because it enables underwater photosynthesis and facilitates quicker recovery after the water recedes. Applying nitrogen (N) after submergence helps to minimize the decrease in chlorophyll a and b levels following de-submergence. Additionally, higher N treatments result in even less decline. The combination of applying phosphorus at the base and nitrogen after flooding helped to maintain higher levels of chlorophyll a and b, non-structural carbohydrates, and photosynthetic rate in plants immersed with clear and turbid water (Gautam *et al*., 2014a). Vergara and Ismail (2006) observed that the introduction of SUB1 resulted in a decrease in the plant’s responsiveness to ethylene, thereby leading to a reduction in chlorophyll degradation while submerged. The intact chlorophyll in the plants boosted their chances of survival by enabling them to produce more energy while submerged and to resume active development after being submerged.

Similar to the variations in total chlorophyll concentrations, the levels of pre-submergence total soluble sugars also differed across all treatments as a result of the varying amounts of basal nitrogen application (as shown in Table 5). However, a notably stronger correlation of 0.84* was found between the concentration of pre-submergence total soluble sugars and the survival rate after 20 days (as indicated in Table 7). Multiple prior investigations have found similar findings, emphasizing the significance of pre-submergence stem polysaccharides in submergence tolerance (Ram *et al*., 2002; Srivastava *et al*., 2007). A significant and robust positive association was found between the survival of the plants after 20 days and the concentration of total soluble sugars that were still present in the stems after being removed from water (correlation coefficient of 0.87*; see Table 7). This suggests that the survival of plants is contingent upon the quantity of remaining carbohydrates in the plant both prior to and following submergence.

### Effect of N supply and application time on survival, post-submergence recovery and yield

The present study examined the impact of pre- and post-submergence nitrogen treatment on specific physiological characteristics linked to tolerance, specifically the overall chlorophyll and soluble sugars content. The survival of plants declined when they were submerged, and this decline was more pronounced when no basal nitrogen was provided. Basal nitrogen also contributed to a decrease in tiller mortality. Application of basal nitrogen appeared to have an impact on the distribution of assimilates, maintaining a balanced ratio between leaves and stems, elevating levels of soluble and insoluble proteins, and ultimately leading to improved survival rates. Table 5 shows that treatments with pre-submergence N application resulted in less significant decreases in both total chlorophyll content and total soluble sugar concentrations. The initial seedling vigor and shoot glucose concentration were considerably improved by applying basal P coupled with N. The study by Gautam *et al*. (2015) found that it improved the chances of survival and regrowth during the period of recovery after being submerged. The use of potassium (K) enhances the ability of plants to withstand submergence by stimulating plant growth and metabolism, as demonstrated by Dwivedi *et al*. (2017). Furthermore, the application of potassium along with basal phosphorus has been found to have a more advantageous impact. This is because the role of phosphorus in mitigating damage caused by submergence has been extensively documented (Gautam *et al*., 2014a). The application of nutrients, specifically nitrogen (N), phosphorus (P), and potassium (K), during submergence and after submergence circumstances, benefits the rice crop by promoting tiller regeneration, maintaining high levels of chlorophyll and total soluble sugar, and enhancing the plants’ ability to withstand stress (Marschner *et al*., 2012; Singh *et al*., 2018). treatment of basal potassium (K) and phosphorus (P), in addition to post-flood nitrogen (N) treatment, dramatically enhanced survival rates by restoring chlorophyll and total soluble sugar levels, hence promoting increased photosynthesis, leaf development, and overall plant growth following submergence (Gautam *et al*., 2014b). The study found significant positive associations between the survival of 5-day-old Arabidopsis seedlings and the levels of total chlorophyll and total soluble sugars before and after submergence. These findings are summarized in Table 7. The rapid and significant increase in crop growth during a time of submergence might be seen as the crop’s re-establishment after being damaged by a flash flood (Ram *et al*, 2009). Pandey (2013) and Mackill *et al*. (2012) suggested that it would be beneficial to apply a tiny additional quantity of nitrogen, ideally one week after the floodwaters had receded. Full submersion of plants leads to an increase in the production of ethylene and its buildup in various plant tissues. This is a significant factor in the degradation of chlorophyll, the aging of leaves, the reduction of stomatal conductance, and the decrease in the photosynthetic rate of submerged plants (Mackill *et al*., 2012; Ismail *et al*., 2012). Singh *et al*. (2014) found that shoot carbohydrates were rapidly depleted during submergence, leading to severe post-oxidative damage after de-submergence. However, the survival of the Swarna-Sub1 cultivar, which had the SUB1 gene introduced, was significantly enhanced by the addition of nitrogen (N), phosphorus (P), and potassium (K). Singh *et al*. (2014) found that the increased regrowth of plants after being brought out of water was linked to the initial application of nitrogen in the field, rather than the nitrogen applied in the nursery along with phosphorus and potassium. Therefore, dividing the application of nitrogen fertilizer into four separate doses (T2; Table 1) offers a chance to enhance the efficiency of nitrogen utilization, decrease the loss of applied inorganic nitrogen, and mitigate yield losses in situations prone to submergence.

The occurrence of submergence typically resulted in a postponement of 50% flowering and maturity. This delay can be attributed to the shedding of leaves and/or shoots during submergence, as well as their partial replacement by new growth during the subsequent recovery period. Following de-submergence, there is a general delay in the process of flowering and maturity. This delay is due to the time required for the surviving plants to recuperate and restart their normal vegetative development. Additionally, the plants need to overcome the damage incurred during and after submergence (Behera *et al*., 2019). Delayed flowering also diminished the duration of grain filling, so directly impacting the grain production. The application of nitrogen in four equal split doses (T2) along with basal phosphorus and potassium resulted in the highest nitrogen use efficiency compared to treatments with no basal nitrogen application or half nitrogen doses as basal (T1 and T3). This is because the continuous availability of nitrogen throughout the growth stages prevented nutrient starvation in the rice plants.

Cassman *et al*. (1994) state that the supply of nitrogen (N) is determined by the presence of N in the soil and the addition of N through fertilizers, as well as the ability of the root system to absorb the available N. Typically, about 30-40% of the total nitrogen fertilizer supplied to flooded rice is effectively used, while the rest remains unused either in the soil or leaches away from it. The yield of cereals is defined by the number of grains per unit of land area, the weight of the grains, and the fraction of grains that are filled.

The grain production of rice in stationary flooded conditions was enhanced by implementing suitable crop and nutrient management practices (Sarangi *et al*., 2015). Bhowmick *et al*. (2014) discovered that administering an extra dose of nitrogen seven days after de-submergence resulted in enhanced survival, post-submergence recovery, yield contributing traits, and overall yield. Gautam *et al*. (2014a and 2015) observed that applying nitrogen to the leaves after submergence, together with phosphorus at the base, led to increased survival, improved management of oxidative damage after submergence, greater yield attributing characteristics, and increased yield in both clear and turbid water submergence conditions with Sub1 cultivars.

## CONCLUSIONS

The study suggests that rice output in rainfed lowlands of Eastern India can be increased by modifying the field procedures employed by farmers, particularly in places that are susceptible to stress. Moreover, modifying the timing, technique, and dosage of nitrogen fertilizer application, in conjunction with the use of tolerant cultivars, has the potential to enhance output. The application of four equal split doses (T2) of nitrogen, together with phosphorus and potassium as basal, improved the rice crop’s ability to withstand submergence. This was achieved by enhancing carbohydrate content, chlorophyll content, and reducing chlorosis, elongation, and lodging. The aforementioned characteristics were manifested in increased rates of survival, crop establishment, and yield of cultivars. The study’s findings demonstrate that applying one-fourth of the N nutrient at the base, along with full doses of P and K, and distributing the remaining N in three equal split doses, is the optimal approach for managing nutrients. This method enhances the resilience of the Sub-1 cultivars, which is particularly advantageous for farmers with limited resources in flood-prone areas of Eastern Uttar Pradesh.

## CRediT Taxonomy

**A. K. Singh:** Principal Investigator, Planning, Methodology, Conceptualization, Supervision. **A.K. Pandey:** Investigation, Data Curation, Writing-original manuscript, review and editing. **Ankit Singh:** Data Collection and Data Analysis. **Deeksha Tiwari:** Data Analysis, Writing-editing. **Bijendra Singh:** Funding acquisition, Visualization and Supervision.

## Acknowledgement

The research was funded by the Centre of Excellence for Rice project, Uttar Pradesh Government, India and facilitated by Hon’ble Vice Chancellor, Dr. Bijendra Singh, A.N.D.U.A&T, Kumarganj, Ayodhya, India.

**Parsed Citations**

